# Splenic tropism of *Plasmodium vivax* in acute infection and spleen-attenuated systemic inflammation

**DOI:** 10.64898/2026.03.25.714340

**Authors:** Steven Kho, Hasrini Rini, Noy N Kambuaya, Syawal Satria, Freis Candrawati, Putu AI Shanti, King Alexander, Benediktus Andries, Aisah R Amelia, Anjana Rai, Kim A Piera, Agatha M Puspitasari, Ristya Amalia, Pak Prayoga, Leo Leonardo, Maulina Hafidzah, Theo Situmorang, Dewi S Margayani, Desandra A Rahmayenti, Pengxing Cao, Enny Kenangalem, Leily Trianty, Damian Oyong, Julie A Simpson, Rintis Noviyanti, Pierre A Buffet, Jeanne R Poespoprodjo, Nicholas M Anstey

## Abstract

**Background:** In chronic asymptomatic *Plasmodium vivax* infections, the spleen accounts for more than 98% of total-body parasite biomass. Whether this splenic tropism also exists in acute infection and how the spleen influences pathogenesis have not been systematically explored.

**Materials and Methods:** In Papua, Indonesia, we compared plasma levels of *P. vivax* lactate dehydrogenase [PvLDH]) and circulating parasitemia in 24 spleen-intact and 25 previously splenectomized patients with acute uncomplicated vivax malaria. Clinical and hematology data were collected and plasma markers of intravascular hemolysis (cell-free hemoglobin [CFHb]), endothelial activation (angiopoietin-2), inflammation (interleukin [IL]-1 beta, IL-6, IL-18, IL-10, tumor necrosis factor-alpha) and neutrophil activation (elastase) were measured by ELISA. Giemsa-based histology in one spleen from an untreated patient splenectomized for trauma during an episode of acute vivax malaria enabled direct assessment of splenic and circulating parasitemia and biomass microscopically.

**Results:** Circulating parasitemia was 4-times higher in splenectomized compared to spleen-intact patients (median 21,100 vs 4,820 parasites/µL, p=0.0002) but total-body *P. vivax* biomass (PvLDH) was 3-times lower in patients without a spleen (median 721 vs 2,140 ng/mL, p=0.026). Parasite staging and greater organ-specific symptoms suggest redistribution of parasites in the absence of a spleen. Linear regression modeling, adjusting for circulating parasitemia, patient age, sex and duration of fever, demonstrated an 8.1-fold higher PvLDH concentration in spleen-intact patients (95% confidence interval [CI]: 3.4-19.5-fold, p<0.0001), indicating a splenic biomass accounting for 89% (95%CI: 77.3-95.1%) of total-body parasites. Histopathology revealed a spleen-to-blood biomass ratio of 10.7, in-line with the PvLDH-based estimate. In spleen-intact patients, splenic *P. vivax* biomass correlated strongly with markers of disease intensity, endothelial activation and systemic inflammation, whereas circulating parasitemia correlated weakly or not at all. Compared to spleen-intact patients, CFHb, endothelial activation and systemic inflammation were higher in splenectomized patients while inflammasome-dependent responses were lower.

**Conclusions:** *P. vivax* is predominantly an infection of the spleen, even in acute clinical vivax malaria. We conservatively estimate that 89% of total-body parasite biomass in acute infection is splenic. While the size of this hidden population correlates with disease intensity, the spleen likely regulates inflammatory pathways and heme-associated pathology.

## Introduction

*Plasmodium vivax* remains the most geographically widespread cause of human malaria, affecting nearly half of the world’s population and accounting for substantial morbidity outside of Africa [1, 2]. Once considered a relatively benign infection [3], vivax malaria is now widely recognised to cause severe disease, recurrent episodes of febrile infections, acute and chronic anemia, thrombocytopenia, adverse pregnancy outcomes and associated mortality [4–7]. The weak association between peripheral blood parasitemia and clinical severity in vivax malaria has led to the notion that circulating parasite density substantially underestimates total-body *P. vivax* biomass, yet the anatomical distribution and pathological significance of non-circulating *P. vivax* parasites in acute clinical infections remains incompletely defined.

Biomarker based studies in uncomplicated and severe vivax malaria have identified the presence of a significant *P. vivax* biomass associated with disease severity [8], greater host perturbations [9] and systemic inflammatory pathology [8]. Histopathology data from historical autopsy series and case reports have reported the presence of intraerythrocytic *P. vivax* parasites in the spleen and bone marrow in clinical disease [10–14]. More recently, analysis of spleens from asymptomatic individuals undergoing splenectomy in Papua, Indonesia, has revealed a large splenic asexual-stage reservoir more than 80-fold greater than circulating biomass, accounting for more than 98% of total-body *P. vivax* biomass [15, 16]. These findings redefine *P. vivax* infection in chronic carriers as an infection developing primarily in the spleen and raises the possibility that the spleen may also serve as the major parasite reservoir in acute clinical vivax malaria.

Direct quantification of parasites in human tissues during untreated acute infections is rarely feasible, limiting the assessment of the relative contribution of splenic versus circulating parasites to total-body *P. vivax* biomass and disease pathogenesis in acute disease. As an alternative, analysis following splenectomy provides a unique model to interrogate spleen pathophysiology and function in human disease. Splenectomized patients are at greater risk of vivax malaria [17] and display greater circulating parasitemia [18], although the impact of spleen removal on total parasite biomass and host responses has not been examined. Plasma *P. vivax* lactate dehydrogenase (PvLDH) has emerged as a robust biomarker of parasite biomass and accounts for parasites from both circulating and non-circulating compartments [8]. Comparison of splenectomized and spleen-intact patients with vivax malaria offers a powerful approach to study splenic pathobiology that can be extrapolated to estimate circulating versus splenic parasite burden.

In this study, we compared plasma PvLDH in spleen-intact and splenectomized patients with acute uncomplicated vivax malaria to estimate the relative contribution of splenic biomass to total-body *P. vivax* burden. This was complemented by direct histopathological quantification of parasites in splenic tissue from an untreated patient. By integrating clinical data, biomarker analysis and cytokine profiling, we further determined the extent to which splenic biomass may contribute to clinical illness and pathogenesis of acute vivax malaria, and how removal of the spleen may reshape inflammatory pathways and disease mechanisms.

## Materials and Methods

### Study site, patients and samples

Timika is a town in the Mimika District of Southern Papua, Indonesia, a lowland partially deforested region with high and perennial malaria transmission. Approximately 37% of the population are infected with *Plasmodium* when assessed by polymerase chain reaction (PCR), with nearly half of infections due to *P. vivax* [19]. Clinical illness with acute uncomplicated malaria is widespread in Timika. In this study, we enrolled two patient groups with acute uncomplicated vivax malaria aged >12 years, comprising of 1) patients with an intact spleen, and 2) patients without a spleen because of previous splenectomy. Acute uncomplicated vivax malaria was defined as microscopy-positive PCR-confirmed *P. vivax* monoinfection with fever (>37.5°C) or history of fever in the past 48 hours, absence of WHO severe malaria criteria [20], and no alternative clinical explanation for their illness. Spleen-intact patients were enrolled as part of a therapeutic efficacy study of dihydroartemisinin-piperaquine between May-December 2023 [21]. Splenectomized patients were identified through a longitudinal prospective cohort of patients undergoing splenectomy in Timika [16], and were enrolled at the time of their first febrile episode of vivax malaria after splenectomy (February 2020 – October 2023). Peripheral venous blood samples were collected prior to antimalarial therapy administered at the site of collection. Automated blood counts were obtained on a Sysmex XP-100 analyzer (Hyogo, Japan). Blood smears were prepared, and samples were frozen as plasma and packed red cells for ELISA and PCR, respectively. Demographic and clinical data were recorded, including clinical assessment of spleen size by Hackett grading.

### Microscopy and PCR

Thick and thin peripheral blood smears were stained with Giemsa and examined by a WHO-certified expert microscopist for *P. vivax* parasitemia, quantified in the thick smear as the number of parasites per 200 white blood cells, or in the thin smear as parasites per 1000 red blood cells. Parasite counts per microliter of blood were calculated based on automated white cell counts or hematocrit data, respectively. The proportions of *P. vivax* developmental stages (rings, trophozoites, schizonts, gametocytes) in peripheral blood were microscopically quantified and expressed as percentages. *P. vivax* monoinfection was confirmed by PCR on genomic DNA extracted from frozen packed red blood cells using the QIAamp DNA Mini kit (Qiagen, Germany). In the splenectomized group, PCR was performed using the AbTES™ Malaria 5 qPCR II kit (AITbiotech, Singapore) according to manufacturer’s instructions. PCR in the spleen-intact group was performed according to methods described previously [21, 22].

### Enzyme-linked Immunosorbent Assays (ELISAs)

Plasma concentration of *P. vivax* lactate dehydrogenase (PvLDH), a measure of total-body *P. vivax* biomass, was determined using an inhouse ELISA as described previously [8, 23]. Plasma levels of the endothelial activation marker, angiopoietin-2 (ang-2), and the pro-inflammatory cytokine, interleukin-6 (IL-6), were determined using the Quantikine™ ELISA Human Ang-2 and Human IL-6 Immunoassays from R&D Systems (Bio-Techne, US). Three other pro-inflammatory cytokines were also measured comprising tumor necrosis factor-alpha (TNF-alpha), interleukin-1 beta (IL-1b) and interleukin-18 (IL-18), as well as the anti-inflammatory marker interleukin-10 (IL-10), all using SimpleStep ELISA® or Human ELISA kits from Abcam (Cambridge, UK). Plasma neutrophil elastase concentrations were determined using the Human Polymorphonuclear Elastase ELISA kit, also from Abcam. Plasma levels of cell-free hemoglobin (CFHb) were measured using the Human Hemoglobin ELISA Quantitation Set (Bethyl Laboratories, Montgomery, TX). Due to artefactual hemolysis from venipuncture in the main spleen-intact patient (cohort 1), CFHb data in splenectomized patients were compared to data available from a separate cohort of spleen-intact patients with acute uncomplicated vivax malaria (cohort 2) [24]. All ELISAs were performed according to manufacturer’s instructions using frozen single-spun heparinized plasma samples (neat or diluted up to 1:10,000) and read on a GloMax plate reader (Promega, Wisconsin, US).

### Spleen tissue histology

Direct histological counting of parasites was performed on one spleen from an untreated patient with incidental acute uncomplicated vivax malaria at the time of major trauma, enrolled as part of our ongoing prospective splenectomy cohort in Timika. Spleen tissue and peripheral blood samples were collected and processed according to procedures described elsewhere [16]. Formalin-fixed paraffin embedded spleen sections were stained with Giemsa and examined by two expert microscopists to quantify splenic parasitemia and total splenic *P. vivax* biomass as described [16], including categorizing *P. vivax* developmental stages and architectural localization (cords, sinus lumens, perifollicular zones, white pulp). Circulating parasitemia was quantified on Giemsa-stained peripheral blood smears, and total circulating *P. vivax* biomass was calculated based on total-body blood volume (estimated using Nadler’s method [16]). Microscopy images were captured on an AxioCam 712c camera coupled to an AxioScope A1 microscope, and annotated using ZEN Software (all Carl Zeiss, Germany). Minor adjustments to brightness/contrast were applied to the entirety of images which did not modify original features.

### Statistical Analysis

All data were analysed using Graphpad Prism v10 (GraphPad, California) or Stata v17 (StataCorp, Texas). The Mann-Whitney U test was used for comparison of continuous variables between spleen-intact versus splenectomized patient, including respiratory rate; red cell parameters (hemoglobin, haematocrit, red cell counts, MCV, MCHC); circulating white cell, platelet and neutrophil counts; circulating parasitemia and the proportion of parasite developmental stages; and plasma levels of CFHb, IL-6, TNF-alpha, ang-2, IL-1b, IL-18, IL-10 and neutrophil elastase. Initially, the distribution of baseline circulating parasitaemia and PvLDH were compared between spleen-intact and splenectomized patients using the Mann-Whitney U test. Estimation of PvLDH-based parasite biomass in the spleen was performed using a linear regression model controlling for circulating parasitemia, patient age, sex and duration of fever (parasitemia and PvLDH values log transformed). Assumptions on PvLDH dynamics governing production and release into plasma from parasites in different compartments are described in the results section. Associations between parasite biomass/parasitemia and markers of disease intensity, inflammation, endothelial activation, hemolysis and neutrophil activation were estimated separately in the spleen-intact and splenectomized groups using the non-parametric Spearman correlation.

### Ethical Approval

This work was approved by the Human Research Ethics Committees of Universitas Gadjah Mada (KE-FK-0505-EC-2019; KE-FK-0445-EC-2023) and Menzies School of Health Research (HREC 2010-1397; 2020-3652). Written informed consent was obtained from all participants.

## Results

### Patient characteristics

In total 49 febrile patients experiencing acute uncomplicated malaria in Timika with PCR-confirmed *P. vivax* monoinfection were included in the study, of which 24 had an intact spleen (cohort 1) and 25 were splenectomized (**Table 1**). Patients without a spleen presented a median of 82 days after splenectomy (interquartile range [IQR]:67-155 days, range:46-522 days). There were no differences in age (median 25 vs 24 years), sex (75% vs 84% males) or the number of days of fever prior to presentation (median 3 vs 2 days) between spleen-intact and splenectomized patients (**Table 1**). There was twice the proportion of Papuans in the spleen-intact group (40% vs 21%; **Table 1**). Compared to spleen-intact patients, there was a significant increase in the respiratory rate in splenectomized patients (median 20 vs 22 breaths per min, p=0.004), and an increase in the proportion with cough (8% vs 24%; **Table 1**). The spleen was palpable in 46% of spleen-intact patients. Hematocrit and circulating red cell, white cell and platelet counts, but not hemoglobin, were significantly reduced in spleen-intact compared to splenectomized patients (**Table 2**).

**Table 1.** Baseline characteristics of patients with acute uncomplicated vivax malaria.

|  | Spleen-intact<br>Cohort 1<br>n=24 | Splenectomized<br>n=25 |
| --- | --- | --- |
| Age in years<br>(median [IQR]) | 25 (20.3-36.3) | 24 (18-35.5) |
| Sex<br>(n of males, [%]) | 18 (75) | 21 (84) |
| Ethnicity<br>(n of Papuans, [%]) | 5 (21) | 10 (40) |
| Body temp. in °C<br>(median [IQR]) | 36.8 (36.6-37.2) | 37.1 (36.5-38.5) |
| Fever days<br>(median [IQR]) | 3 (2-3) | 2 (2-3.5) |
| Cough<br>(n [%]) | 2 (8) | 6 (24) |
| Respiratory rate per min<br>(median [IQR]) | 20 (20-22) | 22 (20-24) |

Abbreviations: IQR, interquartile range.

**Table 2.**
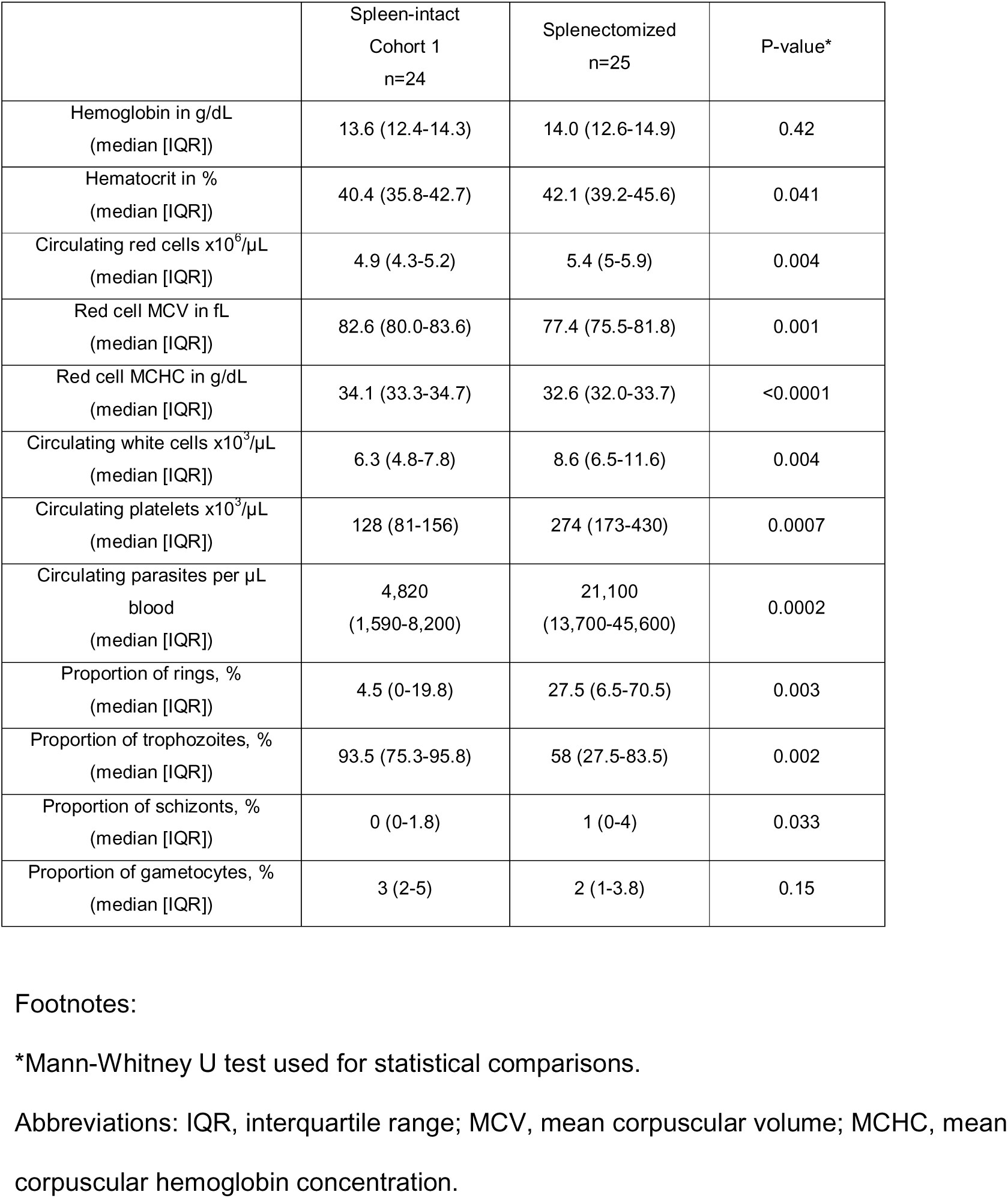
Laboratory and microscopy data in patients with acute uncomplicated vivax malaria.

### Parasitemia in circulating blood is increased in patients without a spleen

The median circulating parasitemia in patients without a spleen was 21,100 parasites per µL (IQR:13,700-45,600 per µL), four times higher than the circulating parasitemia in spleen-intact patients of 4,820 parasites per µL (IQR:1,590-8,200 per µL, p=0.0002; **Table 2**; **Figure 1A**). Compared to spleen-intact patients, circulating blood in splenectomized patients had a significantly higher proportion of ring stages (median 27.5% vs 4.5%, p=0.003) and schizonts (median 1% vs 0%, p=0.033), and a reduction in trophozoite stages (median 58% vs 93.5%, p=0.002; **Table 2**), consistent with redistribution of asexual *P. vivax* stages in the absence of a spleen. There was no significant difference in the proportion of gametocytes between the two patient groups (median 2% vs 3%, p=0.15; **Table 2**).

**Figure 1.**
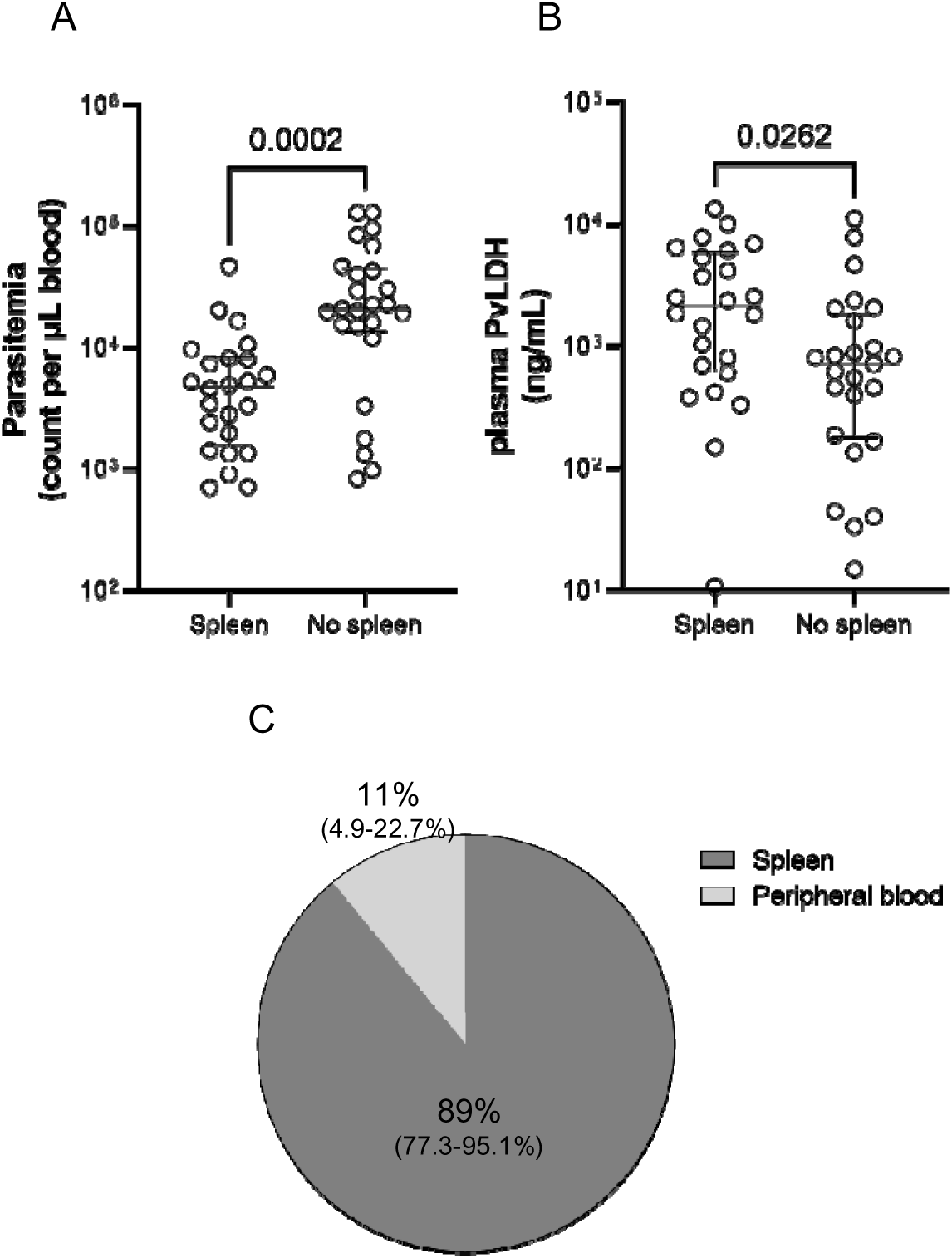
Comparison of A) circulating parasitemia and B) plasma PvLDH concentration in spleen-intact (spleen, n=24, cohort 1) and splenectomized individuals (No spleen, n=25) with acute uncomplicated vivax malaria in Timika, Indonesia. Plasma PvLDH concentration was used as a marker of total-body *P. vivax* biomass. Data points are individual patients with median and interquartile range shown. The Mann-Whitney U test was used for statistical comparison. C) Distribution of *P. vivax* biomass in the spleen and peripheral blood expressed as percentages and 95%CI, determined based PvLDH-based regression model estimates. Abbreviations: PvLDH, *Plasmodium vivax* lactate dehydrogenase; 95%CI, 95% confidence interval.

### Total-body *P. vivax* biomass is greater in spleen-intact than splenectomized patients

Median plasma PvLDH concentration in patients with a spleen was 2,140 ng/mL (IQR:630-5,920 ng/mL), three times higher than the PvLDH concentration in splenectomized patients of 721 ng/mL (IQR:179-1,840 ng/mL, p=0.026; **Figure 1B**). These data are consistent with the presence of a large non-circulating *P. vivax* biomass in patients with a spleen that is either absent or considerably reduced in splenectomized patients.

### The majority of parasite biomass in acute uncomplicated vivax malaria is splenic

The relative size of splenic *P. vivax* biomass was calculated based on the principal assumption that the difference in plasma PvLDH in patients with and without a spleen represents the amount of PvLDH generated and released by the splenic reservoir. Two other conservative assumptions were also made: 1) plasma PvLDH in patients without a spleen represents PvLDH generated and released only by circulating *P. vivax* parasites, and 2) the amount of PvLDH per parasite entering plasma is equivalent between splenic and circulating parasites. Using a linear regression model controlling for circulating parasitemia, patient age, sex and duration of fever, our data indicate that patients with a spleen versus those without a spleen have an 8.1-fold increase in plasma PvLDH concentration (95%CI:3.4-19.5-fold, p<0.0001). Thus, considering the aforementioned assumptions, this indicates that the size of *P. vivax* biomass in the spleen is conservatively 8.1 times (95%CI: 3.4-19.5 times) higher than circulating biomass, with splenic biomass therefore estimated to account for 89% (95%CI: 77.3-95.1%) of total *P. vivax* biomass in acute uncomplicated vivax malaria (**Figure 1C**).

### Confirmation of biomass ratio by splenic histology

An adolescent Papuan male with acute uncomplicated vivax malaria underwent splenectomy due to splenic injury from trauma prior to commencement of antimalarial therapy (**Supplementary Table 1**), enabling direct Giemsa-based estimation of the spleen-to-blood parasitemia and biomass ratios by two trained microscopists (**Figure 2A**). Non-phagocytosed *P. vivax-*infected erythrocytes of all asexual-developmental stages were observed in the spleen (**Figure 2B**). Asexual parasitemia in the spleen was concordant between microscopists (2.9% vs 2.3%), with results from the first reader reported below. Circulating parasitemia in this patient was 238 parasites/µL blood and the spleen-to-blood parasitemia ratio was 570. Calculations incorporating total blood volume and spleen weight estimated a total of 14.3×10^9^ parasites in the spleen and 1.34 ×10^9^ parasites in circulating blood. When calculating the spleen-to-blood biomass ratio in this patient, *P. vivax* biomass in the spleen was 10.7 times higher than biomass in circulating blood, similar to our PvLDH-based estimate.

**Figure 2.**
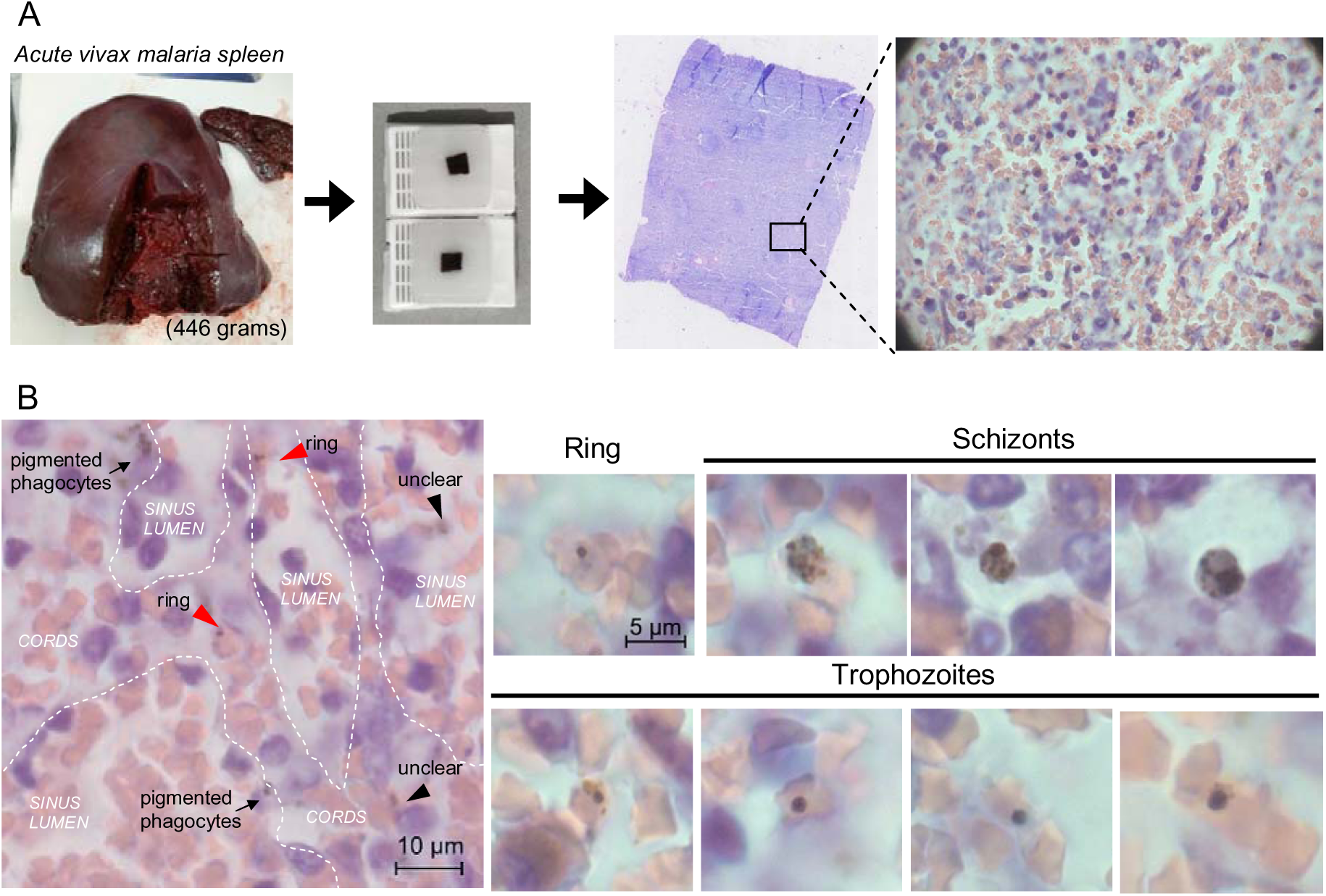
Histopathological spleen assessment in a single patient with untreated acute uncomplicated vivax malaria. (A) Formalin-fixed paraffin embedded spleen tissue blocks were prepared, sliced to 5 µm in thickness and stained with Giemsa for parasite counting and staging by two trained microscopists. (B) Intact non-phagocytosed *P. vivax-*infected erythrocytes of all asexual developmental stages (rings, trophozoites, schizonts) were observed in different architectural zones including the red pulp cords and sinus lumens (representative images shown). Parasites were counted on a Carl Zeiss AxioScope A1 microscope and images captured on an AxioCam 712c, annotated using ZEN Software.

Within the spleen, asexual stages were localized predominantly in the red-pulp cords (92.2%) and a minority in the sinus lumens (6.6%) and perifollicular zones (1.2%). The parasitemia in these regions were 7.3%, 0.3% and 0.5%, respectively. Within the red-pulp, asexual stages comprised 48.3% rings, 20% trophozoites and 31.7% schizonts. In the perifollicular zones, asexual stages comprised 66.7% rings and 33.3% trophozoites. No parasites were observed in the non-circulatory spaces of the white-pulp, and all parasites found in circulating blood were of trophozoite stage.

### In spleen-intact patients, hidden splenic biomass correlates with markers of illness, endothelial activation and systemic inflammation

Correlations between parasite biomass and markers of illness, inflammation, endothelial activation and neutrophil activation were examined in spleen-intact patients to determine the extent to which *P. vivax* biomass (in the spleen) contributes to pathogenesis of vivax malaria (**Table 3**). Overall, increased splenic *P. vivax* biomass was associated with greater disease intensity, correlating significantly in spleen-intact patients with the levels of leucocytosis (r=0.62, p=0.001), ang-2 (endothelial activation) (r=0.44, p=0.033) and key systemic pro-inflammatory markers IL-6 (r=0.45, p=0.02) and TNF-alpha (r=0.45, p=0.042; **Table 3**). Trends towards correlations were also observed between splenic *P. vivax* biomass and duration of fever (r=0.37, p=0.072), the level of thrombocytopenia (r=-0.37, p=0.074) and plasma concentrations of the anti-inflammatory cytokine IL-10 (r=0.37, p=0.090; **Table 3**).

**Table 3.** Spearman correlation coefficients of circulating parasitemia and total-body *P. vivax* biomass with markers of illness, inflammation, endothelial activation and neutrophil activation.

|  | Spleen-intact (Cohort 1) |  |  |  |  | Splenectomised |  |  |  |  |
| --- | --- | --- | --- | --- | --- | --- | --- | --- | --- | --- |
|  | n | Circulating parasitemia | P-value | Total biomass (PvLDH) | P-value | n | Circulating parasitemia | P-value | Total biomass (PvLDH) | P-value |
| Fever days | 24 | 0.29 | 0.17 | 0.37 | 0.072 | 25 | -0.22 | 0.29 | 0.35 | 0.090 |
| WBC count | 24 | 0.49 | 0.016 | 0.62 | 0.001 | 25 | -0.36 | 0.078 | -0.24 | 0.26 |
| Platelet count | 24 | -0.14 | 0.51 | -0.37 | 0.074 | 25 | -0.02 | 0.93 | -0.53 | 0.006 |
| Ang-2 | 24 | 0.22 | 0.31 | 0.44 | 0.033 | 25 | 0.25 | 0.23 | 0.55 | 0.005 |
| IL-6 | 24 | 0.52 | 0.008 | 0.45 | 0.028 | 25 | 0.60 | 0.002 | 0.65 | 0.0004 |
| TNF-alpha | 21 | 0.13 | 0.58 | 0.45 | 0.042 | 23 | 0.61 | 0.002 | 0.45 | 0.031 |
| IL-1b | 24 | -0.16 | 0.45 | 0.16 | 0.45 | 18 | -0.17 | 0.51 | 0.14 | 0.58 |
| IL-18 | 24 | 0.17 | 0.43 | 0.26 | 0.22 | 24 | 0.37 | 0.076 | 0.01 | 0.99 |
| IL-10 | 22 | 0.35 | 0.11 | 0.37 | 0.090 | 23 | 0.23 | 0.28 | 0.51 | 0.013 |
| NE | 20 | 0.09 | 0.70 | 0.14 | 0.54 | 25 | 0.01 | 0.99 | 0.23 | 0.27 |

The following number of patients had plasma measurement below the limit of detection of the assays and were therefore excluded from correlations: 3 TNF-alpha and 2 IL-10 in the spleen-intact group; 2 TNF-alpha, 7 IL-1b, 1 IL-18 and 2 IL-10 in the splenectomized group. Measurement of NE in 4 spleen-intact patients were above the upper limit of detection of the assay and were also excluded.
Abbreviations: WBC, white blood cell; Ang-2, Angiopoietin-2; IL-6, interleukin-6; TNF-alpha, tumour necrosis factor-alpha; IL-1b, interleukin-1 beta; IL-18, interleukin-18; IL-10, interleukin-10; NE, neutrophil elastase; PvLDH, *Plasmodium vivax* lactate dehydrogenase

In contrast to the correlations with splenic *P. vivax* biomass, circulating *P. vivax* parasitemia in spleen-intact patients was only associated with greater plasma IL-6 levels (r=0.52, p=0.008) and white cell counts (r=0.49, p=0.016), and was not associated with any of the other markers that were tested (**Table 3**), consistent with splenic *P. vivax* biomass contributing to inflammatory pathogenesis to a greater degree than circulating parasites in vivax malaria. Plasma levels of neutrophil activation (elastase) and inflammasome-dependent inflammatory cytokines IL-1b and IL-18 did not correlate with splenic *P. vivax* biomass nor circulating parasitemia in the spleen-intact group (**Table 3**).

### Plasma levels of cell-free hemoglobin are high in splenectomized patients with acute vivax malaria

Cell-free hemoglobin, a toxic and major trigger for inflammation [25, 26], was measured in the plasma of splenectomized patients and compared to CFHb data available in a separate historical group of spleen-intact patients with acute uncomplicated vivax malaria (cohort 2 (n=36); baseline characteristics in **Supplementary Table 2**). In splenectomized patients, CFHb levels were more than twice the levels observed in spleen-intact patients (median 73,000 vs 30,400 ng/mL, p=0.0001; **Figure 3A**). In these splenectomized patients, increased plasma CFHb levels was associated with greater *P. vivax* biomass (r=0.50, p=0.011), longer duration of fever (r=0.59, p=0.002), increased respiratory rate (r=0.40, p=0.046), thrombocytopenia (r=-0.64, p=0.001), endothelial activation (ang-2) (r=0.40, p=0.05) and systemic inflammation (IL-6) (r=0.42, p=0.038; **Table 4**). There was also a trend towards higher body temperature (r=0.37, p=0.073; **Table 4**). In contrast to these observations in splenectomized patients, no associations between CFHb levels and markers of inflammation were present in spleen-intact patients (**Table 4**). Interestingly, pigmented phagocytes were histologically observed in the single spleen from an acute infection (**Figure 2B**), in-line with processes related to hemozoin engulfment or scavenging of free hemoglobin by macrophages or other cells in the spleen [27–29]. Taken together, these data are consistent with a splenic role in removing/attenuating CFHb levels and potentially other toxic by- products from the circulation through innate sensing, a key role that is lost in splenectomized patients leading to circulating heme-mediated toxicity and pathogenesis.

**Figure 3.**
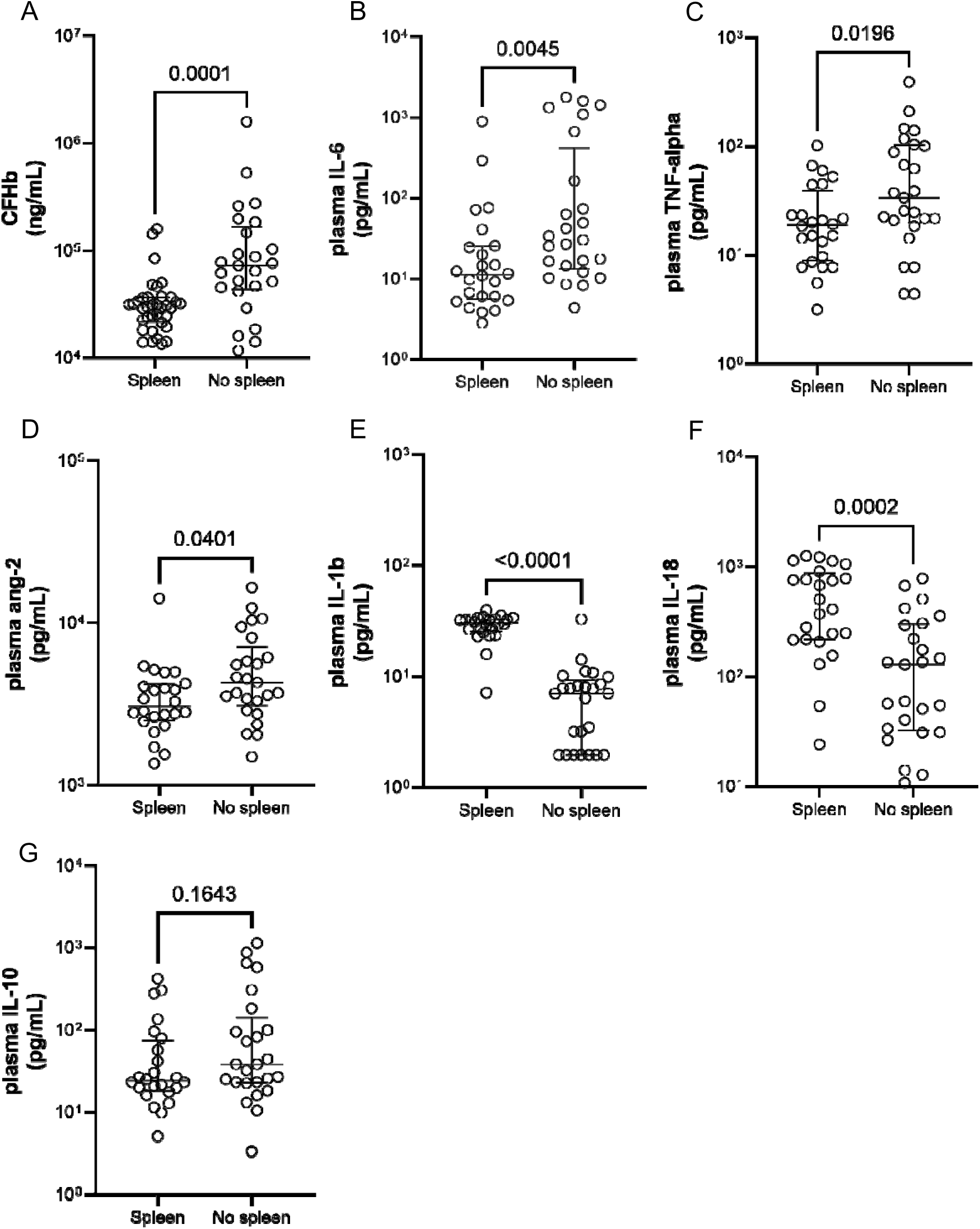
Comparison of plasma concentrations of A) CFHb, B) IL-6, C) TNF-alpha, D) ang-2, E) IL1-beta, F) IL-18 and G) IL-10 in spleen-intact (spleen) and splenectomized individuals (No spleen, n=25) with acute uncomplicated vivax malaria in Timika, Indonesia. Data points are individual patients with median and interquartile range shown. All measurements in the spleen-intact group were from cohort 1 (n=24), except CFHb (cohort 2, n=36). Measurements below the detection limit of each marker were assigned a value of half the lower limit of detection of the assay, comprising of the following number of patients: 3 TNF-alpha and 2 IL-10 in the spleen-intact group, and 2 TNF-alpha, 7 IL-1b, 1 IL-18 and 2 IL-10 in the splenectomized group. The Mann-Whitney U test was used for statistical comparison. Abbreviations: CFHb, cell-free hemoglobin; IL-6, interleukin-6, TNF-alpha, tumour necrosis factor-alpha; ang-2, angiopoietin-2; IL-1b, interleukin-1 beta; IL-18, interleukin-18; IL-10, interleukin-10.

**Table 4.**
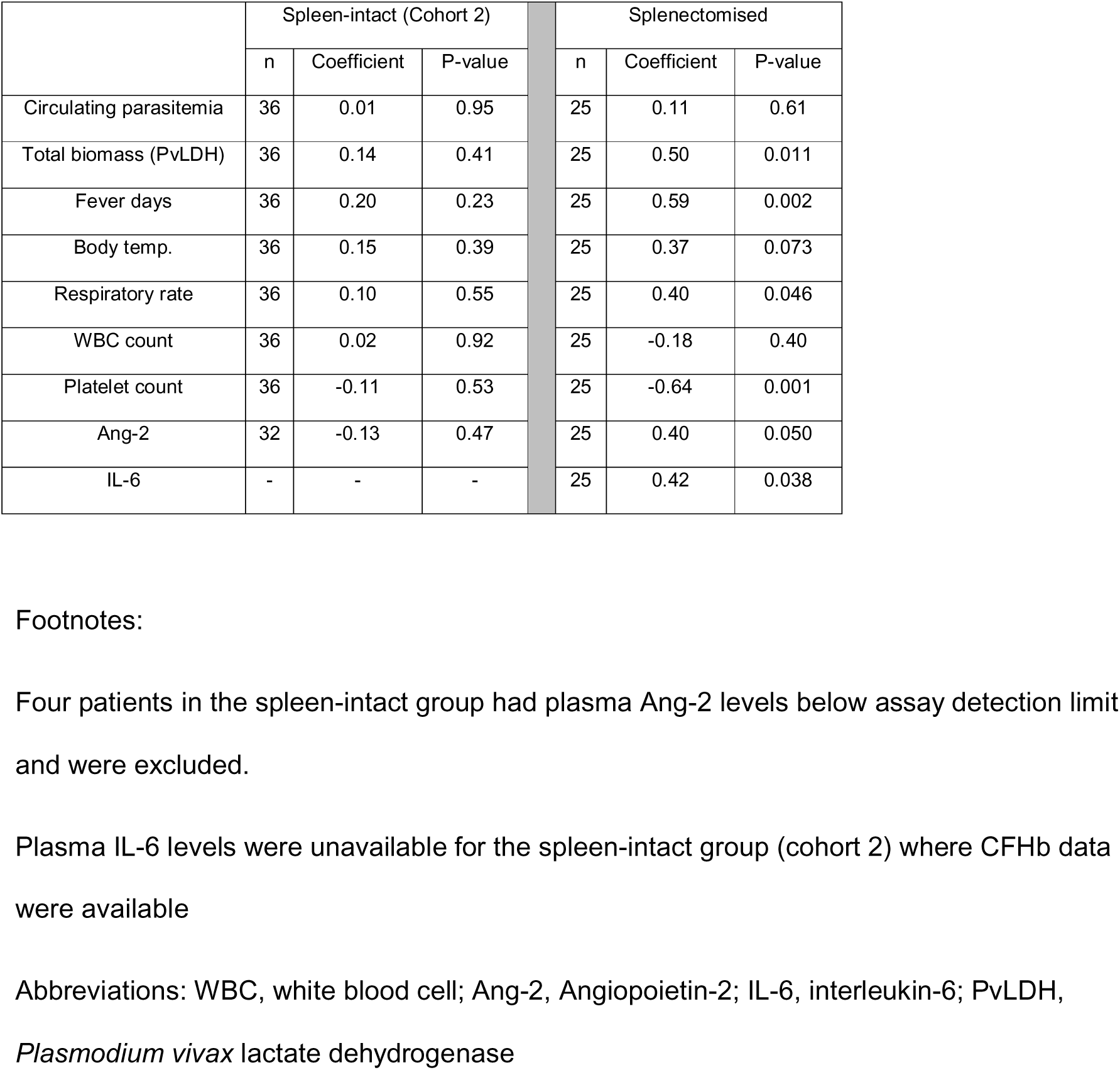
Spearman correlation coefficients of cell-free hemoglobin with circulating parasitemia, total-body *P. vivax* biomass, endothelial activation and markers of illness.

|  | Spleen-intact (Cohort 2) |  |  | Splenectomised |  |  |
| --- | --- | --- | --- | --- | --- | --- |
|  | n | Coefficient | P-value | n | Coefficient | P-value |
| Circulating parasitemia | 36 | 0.01 | 0.95 | 25 | 0.11 | 0.61 |
| Total biomass (PvLDH) | 36 | 0.14 | 0.41 | 25 | 0.50 | 0.011 |
| Fever days | 36 | 0.20 | 0.23 | 25 | 0.59 | 0.002 |
| Body temp. | 36 | 0.15 | 0.39 | 25 | 0.37 | 0.073 |
| Respiratory rate | 36 | 0.10 | 0.55 | 25 | 0.40 | 0.046 |
| WBC count | 36 | 0.02 | 0.92 | 25 | -0.18 | 0.40 |
| Platelet count | 36 | -0.11 | 0.53 | 25 | -0.64 | 0.001 |
| Ang-2 | 32 | -0.13 | 0.47 | 25 | 0.40 | 0.050 |
| IL-6 | - | - | - | 25 | 0.42 | 0.038 |

Four patients in the spleen-intact group had plasma Ang-2 levels below assay detection limit and were excluded.
Plasma IL-6 levels were unavailable for the spleen-intact group (cohort 2) where CFHb data were available
Abbreviations: WBC, white blood cell; Ang-2, Angiopoietin-2; IL-6, interleukin-6; PvLDH, *Plasmodium vivax* lactate dehydrogenase

### Markers of systemic inflammation and endothelial activation are elevated in splenectomized patients, in contrast with reduced inflammasome-dependent cytokines

Compared to spleen-intact patients, splenectomized patients displayed markedly elevated plasma levels of pro-inflammatory cytokines IL-6 (11.4 vs 30.5 pg/mL, p=0.0045; **Figure 3B**) and TNF-alpha (19.0 vs 34.0 pg/mL, p=0.020; **Figure 3C**), as well as a small but significant increase in plasma ang-2 (3,080 vs 4,320 pg/mL, p=0.0401; **Figure 3D**), indicating greater systemic inflammation and endothelial activation in the absence of a spleen in vivax malaria. Elevation of these three markers in splenectomized patients correlated significantly with high *P. vivax* biomass (IL-6 r=0.65, p=0.0004; TNF-alpha r=0.45, p=0.031; ang-2 r=0.55, p=0.005) and circulating parasitemia (IL-6 r=0.60, p=0.002; TNF-alpha r=0.61, p=0.002; **Table 3**). These relationships with circulating parasite load were stronger than those with splenic *P. vivax* biomass in spleen-intact patients (**Table 3**). Collectively, these data are consistent with defective regulation of host inflammatory responses when the spleen is absent, despite lower parasite biomass overall (**Figure 1B**).

In contrast, profiling of inflammasome-dependent cytokines displayed differing patterns, with splenectomized patients producing substantially lower levels of plasma IL-1b (7.1 vs 30.6 pg/mL, p<0.0001; **Figure 3E**) and IL-18 (129 vs 459 pg/mL, p=0.0002; **Figure 3F**) compared to spleen-intact patients. In addition, neither cytokine correlated with *P. vivax* biomass in splenectomized individuals (**Table 3**), as observed in the spleen-intact group.

### Anti-inflammatory and neutrophil responses

Despite greater systemic inflammation in splenectomized patients (**Figure 3B,C**), plasma levels of the anti-inflammatory cytokine IL-10 were similar between patients with and without a spleen (24.4 vs 38.6 pg/mL, p=0.16; **Figure 3G**). The correlation between IL-10 and *P. vivax* biomass was only significant in the splenectomized group (r=51, p=0.013; **Table 3**).

Notably, IL-10 levels strongly correlated with IL-6 in both spleen-intact and splenectomized patients (r=0.86 and r=0.78, both p<0.0001), but only correlated with ang-2 in the splenectomized patients (r=0.62, p=0.001). IL-10 also strongly correlated with increased body temperature in the splenectomised patients (r=0.91, p<0.0001), and only weakly in the spleen-intact patients (r=0.37, p=0.094). Thus, while IL-10 production was responsive to systemic inflammation in both patient groups, our data point towards a defective compensatory anti-inflammatory response being elicited in direct proportion to parasite load, endothelial activation and degree of fever, in the absence of a spleen.

Neutrophil counts were significantly higher in the circulation of splenectomized compared to spleen-intact patients (5,550 vs 4,200 per µL, p=0.026; **Figure S1A**), suggesting potential redistribution to the blood following splenectomy. In contrast, the level of systemic neutrophil activation (plasma NE) was significantly lower in splenectomized compared to spleen-intact patients (675 vs 1,030 ng/mL, p=0.0022; **Figure S1B**), and normalising NE to circulating neutrophil counts did not alter this relationship (p<0.0001; **Figure S1C**). In both patient groups, there was no correlation between NE levels and *P. vivax* biomass, circulating parasitemia (**Table 3**) or any of the other plasma markers (data not shown).

## Discussion

Circulating parasites were long thought to be the only component of total-body *P. vivax* biomass, and therefore the sole trigger of inflammation in acute clinical disease. Cytoadherence-based sequestration of infected erythrocytes in small vessels is much lower than with *P. falciparum* [5, 30]. Here, by comparing PvLDH levels between spleen-intact and splenectomized patients with acute uncomplicated vivax malaria, we show that the human spleen is the dominant reservoir of parasites, accounting for 89% of total-body parasite biomass. Spleen histopathology in one patient confirms this vastly predominant tropism. Clinical correlates and cytokine profiling indicate that splenic parasite biomass is the main driver of disease pathogenesis in acute infections. Along with spleen-related cytopenias, this process points to a pathogenic role for the spleen in acute vivax malaria. Conversely, comparison of hemolysis and other inflammatory markers between groups strongly suggests that the spleen also clears CFHb and controls both systemic inflammation and inflammasome-related pathways. The spleen thus harbours the bulk of the parasite population (that induces inflammation), but also markedly tempers this inflammation and related pathogenic mechanisms.

In patients with a spleen, total-body parasite biomass as measured by plasma PvLDH was over eight-fold greater than biomass attributed conservatively to circulating parasites alone. This large hidden reservoir was effectively absent in splenectomised patients who exhibited higher circulating parasitemia but markedly reduced total-body biomass. The simplest interpretation is therefore that most *P. vivax* parasites reside within splenic tissue rather than in circulation, as confirmed by direct microscopic quantification in splenic sections. This splenic tropism is consistent with our findings in chronic asymptomatic *P. vivax* infections from the same endemic setting [15]. Splenic accumulation was ten times lower in acute infections in the current study than in previously explored chronic carriers, with the spleen-to-blood biomass ratio being 8.1 in acute infections and 81 in chronic carriers [16]. Despite this difference in the magnitude of splenic tropism, histology in acute vivax malaria showed that accumulation predominates in the red pulp cords as also seen in chronic infections [11, 16], and more recently in acute malaria due to *P. ovale* [31].

Elevated circulating parasitemia after splenectomy is consistent with other reports in *P. vivax* [18], *P. falciparum* [32] and *P. knowlesi* malaria [33]. Higher circulating parasitemia and redistribution of asexual *P. vivax* stages in splenectomized patients likely reflects increased presence of parasite stages that would otherwise be at least partly retained in the spleen. Very early *P. vivax* rings that have recently invaded CD71+ reticulocytes (<3 hours) [34], as well as segmenting schizonts [35], have reduced deformability and are therefore susceptible to biomechanical splenic trapping [36–38]. Our data is consistent with this, with rings and schizonts present in the spleen but not blood of the single patient with histology, and their proportions were found to be increased in circulating blood after splenectomy. On splenic sections, the predominance of asexual parasites in the slow-circulating cords, and at much higher parasitemias compared to other splenic zones, is further supportive of biomechanical accumulation. Determining the respective contributions of such mechanical retention versus cytoadherence-based sequestration [39–41] requires further study.

In non-human primates, splenic tropism has also been observed [42]. However, in splenectomized non-human primates there was major redistribution of *P. vivax* load to other organs after spleen removal, particularly to the liver, lungs and bone marrow [42, 43]. *P. vivax* parasites have been reported in these non-splenic tissues in humans [12, 13, 44]. Increased frequency of cough and elevated respiratory rate in splenectomized patients with vivax malaria was notable in our study, consistent with possible redistribution and increased pathology in the lungs, and supportive of murine data identifying the lungs as a major site for biomechanical filtration second only to the spleen [45].

The large splenic biomass in acute vivax malaria has implications for recrudescence of infection following standard blood-stage schizonticidal treatment. Studies in acute falciparum malaria have shown that failure to clear infections with these drug regimens and consequent risk of recrudescent infections is strongly related to pre-treatment total parasite biomass [46, 47]. With the lower circulating parasitemias generally found in acute vivax malaria [5] and the previous assumption (now refuted in this study) that all *P. vivax* parasites in acute infection circulate, it has long been assumed that the initial parasite biomass in acute vivax malaria would be more than adequately cleared by standard schizonticidal treatment regimens. The 8-fold greater biomass of *P. vivax* hidden in the spleen means that total pre-treatment parasite biomass is much greater than previously assumed. Thus, as in falciparum malaria, in those with the highest starting parasite biomass, recurrent vivax infections may arise from recrudescence of asexual parasites incompletely cleared by standard schizonticidal drug regimens, particularly if splenic parasites are less susceptible to treatment [48, 49]. Most recurrent vivax infections are currently attributed to relapsing infections from activation of latent liver stages which are prevented by concurrent 8-aminoquinoline treatments such as primaquine [5]. However, not all 8-aminoquinoline-preventable recurrences may be true relapses from hepatic hypnozoite activation. Primaquine, activated through an enzymatic pathway present in liver, bone marrow and likely spleen [50], may also contribute to killing of asexual stages in the spleen and prevent their release back into the circulation to cause recurrence [51]. The proportion of recurrent *P. vivax* infections that are currently attributed to relapses but may be arising from recrudescence of an inadequately cleared asexual-stage splenic biomass, is unknown.

Our correlative assessment between parasite biomass and markers of disease intensity (fever duration, leucocytosis, thrombocytopenia) and inflammation (IL-6, TNF-alpha, ang-2) suggest that splenic *P. vivax* biomass contributes to a greater extent to acute disease pathogenesis than does the circulating biomass, as reflected in previous studies [8, 9]. The spleen can also contribute to increased disease intensity through hypersplenism causing pancytopenia in individuals with spleens acutely enlarged by congestion [36, 52, 53], as seen here, and possibly also through spleen-mediated mechanisms of cell removal [54]. Beyond its role as a parasite reservoir linked to enhanced inflammatory signals, our study suggests that, conversely, the spleen clears toxic hemolytic by-products during vivax malaria. Splenectomized patients had markedly higher plasma CFHb concentrations, more than twice those observed in spleen-intact patients. In splenectomized patients, CFHb was strongly associated with fever duration, thrombocytopenia, ang-2, IL-6, and parasite biomass. Such relationships were absent in spleen-intact patients, consistent with effective splenic control of free heme. Histological identification supports active splenic uptake of free hemozoin, hemoglobin or heme-containing cell components consistent with the scavenging roles of red-pulp macrophages and endothelial cells [55], with many of them containing pigment [27–29]. In patients with intact spleens, CFHb is associated with disease severity in falciparum malaria [56, 57] but not in vivax malaria [58], consistent with our findings. Endothelial activation in acute *P. vivax* is therefore probably driven by hemolytic by-products [9] only in the absence of a functional spleen.

Our data point towards excessive and dysregulated inflammatory responses occurring in the absence of a spleen during vivax malaria. Splenectomised patients displayed higher plasma IL-6 and TNF-alpha levels, and these responses correlated strongly with *P. vivax* biomass, more so than in spleen-intact patients. Heme induces IL-6 production [59], consistent with the association between IL-6 and CFHb levels in splenectomized patients and possibly contributing to this increased IL-6 response. Despite loss of splenic B cells and monocytes with splenectomy, and therefore partial loss of IL-6 and TNF-alpha production niches [60, 61], their levels remained excessively high in splenectomized patients. We speculate that the bone marrow, liver and lung may be key alternative sites of IL-6 [60, 61] and TNF-alpha production [62, 63] during vivax malaria in the absence of a spleen, likely exacerbated by redistribution of *P. vivax* parasites to these organs following splenectomy [62]. Elevated ang-2 levels and the stronger correlations with parasite biomass in splenectomized patients are consistent with higher circulating parasitemia and likely greater interaction between circulating parasites and the vasculature endothelial cells that produce ang-2. The weaker correlation between splenic *P. vivax* biomass and ang-2 in spleen-intact patients is consistent with the smaller fraction of *P. vivax* parasites found in the sinus lumens of the spleen that allow interaction with endothelial cells in this region.

Our assessment of cytokines of the IL-1 family implicate the spleen as a major site for inflammasome activation during vivax malaria, supporting a speculative model in which splenic *P. vivax* biomass may contribute indirectly to inflammasome-associated inflammation through secondary processes that are spatially restricted to the splenic microenvironment. Genetic associations implicating inflammasome pathways in vivax malaria indeed support a pathogenic role for IL-1b and IL-18 [64]. Plasma levels of these two inflammasome-dependent cytokines were markedly reduced in splenectomized patients, consistent with the spleen driving inflammasome-dependent processes. Inflammasome activation requires conversion of pro-IL-1b and pro-IL-18 into their active forms, a process requiring caspase-1 and NLRP3 activity [65]. This process is influenced by factors beyond parasite biomass alone, including immune complexes [61], hemozoin [66], parasite DNA [61, 66] macrophages and heme [67], all of which are abundant but largely contained within the spleen. Levels of immune complexes promoting inflammasome activation have previously been shown to be unrelated to parasitemia in vivax malaria [61], consistent with the lack of correlation between IL-1 family cytokines and parasitemia or total biomass in our patients, and consistent with inflammasome processes being driven indirectly in the spleen by malaria-associated factors than parasite load per se.

Finally, our plasma neutrophil elastase measurements indicate greater systemic neutrophil activation occurring in spleen-intact than splenectomized patients. The higher circulating neutrophil count after splenectomy is consistent with post-splenectomy redistribution of splenic neutrophil pools into blood [68]. These splenic populations are likely significant contributors to neutrophil activation in spleen-intact patients [69], with malaria-associated factors abundant in the splenic microenvironment likely important for effective neutrophil priming and degranulation in vivax malaria [70, 71]. Loss of splenic neutrophils and insufficient compensation from redistribution into the blood (as suggested by the NE-neutrophil count ratio data) may explain the reduced elastase levels observed in splenectomized patients, possibly exacerbated by altered composition of circulating neutrophils towards defective phenotypes after splenectomy [72].

Our study has several limitations. Firstly, the comparison of spleen-intact and splenectomised patients was observational and potential effects related to immunity or post-splenectomy adaptations over time could not be controlled. Nevertheless, the two groups were broadly comparable at presentation with respect to age, sex, ethnicity and fever duration, and the timing of *P. vivax* acute attacks were largely within the first 6 months after splenectomy, which likely minimized the effect of such confounders. Secondly, CFHb comparisons required use of an additional historical spleen-intact group due to artefactual hemolysis during venesection in the study’s primary spleen-intact group, although baseline characteristics between the two spleen-intact cohorts were largely comparable. Direct quantification of splenic parasite biomass by histology in acute infection was also only possible in one patient due to rarity of acute infections coinciding with trauma in our prospective splenectomy cohort, although concordance of histology data with the PvLDH-based estimates supports generalisability of our results. Thirdly, while PvLDH is a well-validated biomarker of total-body parasite biomass, it remains an indirect measure and is influenced by parasite PvLDH production, release and clearance dynamics [23] that may differ between anatomical compartments. We conservatively assumed equal PvLDH production and release by circulating and splenic parasites, and that plasma PvLDH in splenectomised patients was attributable to circulating parasites alone. Redistribution of parasites to non-splenic tissues after splenectomy likely underestimates the true splenic contribution to total biomass in spleen-intact patients, and therefore, our estimate of 89% of parasite biomass in the spleen in uncomplicated vivax malaria should be considered a minimum estimate. Finally, the cross-sectional nature of our study limits causal inference of the temporal relationships between parasite biomass, splenic function and host responses that we propose. Further work is warranted to confirm our findings. The extent to which our study can be generalized to severe vivax malaria, other transmission settings, different age groups, or mixed *P. vivax* infections, also remains to be determined.

In conclusion, our study provides the strongest human evidence to date supporting a model in which the spleen plays a dominant role in shaping *P. vivax* biology, pathogenesis, and host responses during acute vivax malaria. By demonstrating that circulating parasitemia represents only a small fraction of total-body infection, and that illness is primarily driven by splenic rather than circulating parasite load, our findings fundamentally redefine long-standing assumptions regarding parasite tropism and pathogenesis in vivax malaria. Further to its pathogenic role as a parasite reservoir, the spleen also emerges as a critical immunopathological regulator mediating control of inflammation and limiting endothelial tissue injury. Loss of the spleen results in redistribution of parasites, increased circulating parasitemia, loss of innate protection from inflammatory triggers and heme-induced toxicity, excessive uncontrolled systemic inflammatory responses, increased microvascular pathology and impaired inflammasome-dependent pathways. Collectively, these data highlight the human spleen as a ‘double-edged sword’ with dual roles in vivax malaria. Its recognition as the central reservoir of *P. vivax* infection has important implications for clinical management, diagnostics, surveillance, therapeutic studies of parasite clearance and origin of primaquine-preventable recurrent infections, and the design of future interventions aimed at reducing morbidity and transmission of vivax malaria.

## Acknowledgements

We thank patients and relatives of patients in Indonesia for their participation and support, staff in the laboratory and operating theatre at RSUD hospital, staff at Wania Clinic, and field and lab staff involved in the Timika TES and MAPLE studies. We are grateful to colleagues at the Timika Research Facility for their support. Thank you to Associate Professor Michelle Boyle for her intellectual input into the manuscript.

## Author Contributions

SK, PB and NMA conceived the study; SK, KAP, PC, JAS, RN, PAB and NMA provided methodology; SK, KAP and DAR performed validation; SK, HR, NNK, SS, FC,BA, ARA, AMP, RA, PP, LL, DAR, MH, TS and DSM performed experiments and investigation; SK, PAIS, KA, AR, EK, LT, JAS, RN, PAB, JRP and NMA provided resources; SK, NMA and PAB performed formal analysis and wrote the original draft; AR, PC, DO, EK, JAS and JRP performed review and editing; SK, PP, DAR and DO performed visualization; SK, EK, LT, RN, PAB, JRP and NMA provided supervision; SK, RN, JRP and NMA performed project administration; and SK, JRP and NMA provided funding acquisition.

## Data availability

Data in this manuscript are available from the corresponding author upon reasonable request.

