## Supplementary Files for "Splenic tropism of *Plasmodium vivax* in acute infection and spleen-attenuated systemic inflammation"

**Supplementary Table 1.** Clinical and laboratory characteristics of acute vivax malaria patient with histopathology data

| Demographic data | Age | 17 years |
| --- | --- | --- |
|  | Sex | Male |
|  | Ethnicity | Papuan |
|  | Weight | 111 kg |
|  | Height | 158 cm |
| Clinical | Splenectomy | Spleen rupture grade V (trauma: traffic accident) |
|  | Spleen weight | 446 gr |
|  | Fever | Yes |
|  | Symptom history | Headache and nauseous in last 2 days |
|  | Malaria history | 1 *P. vivax* episode in the past year |
|  | Total-body blood volume | 5.62 L |
| Laboratory | Hemoglobin | 11.4 g/dL |
|  | White cell count | 23.9 x10^3^/uL |
|  | Platelet count | 185 x10^3^/uL |
|  | Spleen white pulp area | 10.7% |
|  | Spleen red pulp area | 86.2% |
| Parasitology | PfHRP2/Pan RDT | Negative |
|  | Plasmodium PCR | *P. vivax* monoinfection |
|  | Peripheral parasitemia | 0.005 (% red cells); 238 parasites per µL blood |
|  | Splenic parasitemia | 2.85 (% red cells) |
|  | Total peripheral biomass | 1.34 x10^9^ parasites |
|  | Total splenic biomass | 14.3 x10^9^ parasites |

Footnotes:

Abbreviations: PfHRP2, *Plasmodium falciparum* histidine-rich protein-2; RDT, rapid diagnostic test; PCR, polymerase chain reaction.

**Supplementary Table 2.** Baseline characteristics and laboratory data in spleen-intact patients from cohort 2 without artefactual hemolysis at venesection

|  | Spleen-intact  (cohort 2)  n=36 | Splenectomized  n=25 | P-value* |
| --- | --- | --- | --- |
| Age in years  (median [IQR]) | 24 (19.5-31.5) | 24 (18-35.5) | - |
| Sex  (n of males, [%]) | 23 (64) | 21 (84) | - |
| Ethnicity  (n of Papuans, [%]) | 34 (94) | 10 (40) | - |
| Body temp. in °C  (median [IQR]) | 37.0 (36.4-37.9) | 37.1 (36.5-38.5) | - |
| Fever days (median [IQR]) | 4 (3-7) | 2 (2-3.5) | - |
| Cough  (n [%]) | 11 (31) | 6 (24) | - |
| Respiratory rate per min  (median [IQR]) | 20 (19-24) | 22 (20-24) | 0.008 |
| Haemoglobin in g/dL  (median [IQR]) | 11.9 (10.5-13.1) | 13.6 (12.4-14.3) | 0.0001 |
| Haematocrit in %  (median [IQR]) | 34.4 (28.8-37.0) | 42.1 (39.2-45.6) | <0.0001 |
| Circulating red cells x10^6^/µL  (median [IQR]) | 4.5 (4.0-4.8) | 5.4 (5-5.9) | <0.0001 |
| Circulating white cells x10^3^/µL  (median [IQR]) | 5.5 (4.9-7.6) | 8.6 (6.5-11.6) | 0.0001 |
| Circulating platelets x10^3^/µL  (median [IQR]) | 73 (53-107) | 274 (173-430) | <0.0001 |
| Circulating parasites per µL blood  (median [IQR]) | 6,680  (2,280-11,900) | 21,100  (13,700-45,600) | 0.001 |

Footnotes:

*Mann-Whitney test used for continuous variables and chi-squared test used for categorical variables

Abbreviations: IQR, interquartile range

B

A

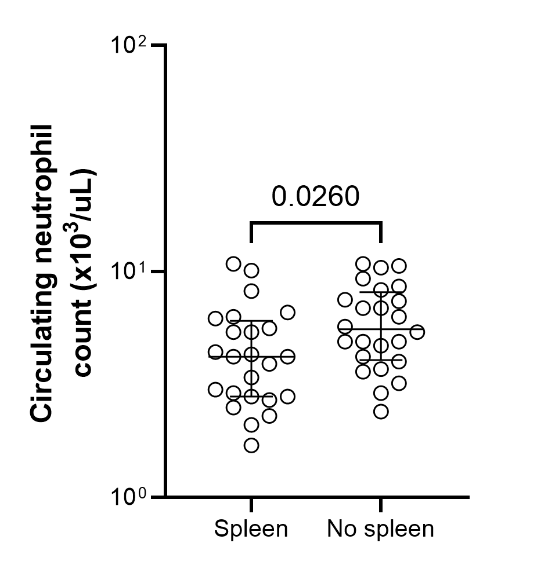

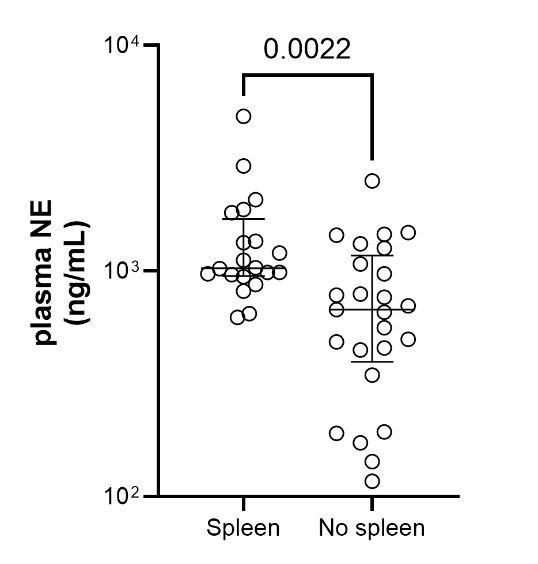

C

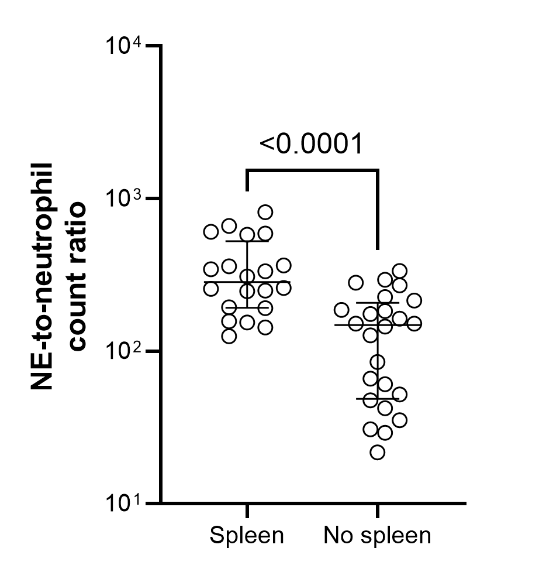

**Supplementary Figure 1.** Comparison of A) circulating neutrophil counts, B) plasma NE concentrations and C) NE-to-neutrophil count ratio in spleen-intact (spleen, cohort 1) and splenectomized individuals (No spleen) with acute uncomplicated vivax malaria in Timika, Indonesia. Data points are individual patients with median and interquartile range shown. Neutrophil counts were available for 24 spleen-intact and 24 splenectomized patients. Plasma NE concentration was used as a marker of systemic neutrophil activation and were available for 20 spleen-intact and 25 splenectomized patients (NE measurements in 4 spleen-intact patients were above the upper limit of detection of the assay and were excluded). The Mann-Whitney test was used for statistical comparison. Abbreviations: NE, neutrophil elastase.
